# The relative contribution of natural landscapes and human-mediated factors on the connectivity of a noxious invasive weed

**DOI:** 10.1101/054122

**Authors:** Diego F. Alvarado-Serrano, Megan Van Etten, Shu-Mei Chang, Regina S. Baucom

## Abstract

Examining how the landscape may influence gene flow is at the forefront of understanding population differentiation and adaptation. Such understanding is crucial in light of ongoing environmental changes and the elevated risk of ecosystems alteration. In particular, knowledge of how humans may influence the structure of populations is imperative to allow for informed decisions in management and conservation as well as to gain a better understanding of anthropogenic impacts on the interplay between gene flow, genetic drift and selection. Here we use genome-wide molecular markers to characterize the population genetic structure and connectivity of *Ipomoea purpurea*, a noxious invasive weed. We likewise assess the interaction between natural and human-driven influences on genetic differentiation among populations. Our analyses find that human population density is an important predictor of pairwise population differentiation, suggesting that the agricultural and/or horticultural trade may be involved in maintaining some level of connectivity across distant agricultural fields. Climatic variation appears as an additional predictor of genetic connectivity in this species. We discuss the implications of these results and highlight future research needed to disentangle the mechanistic processes underlying population connectivity of weeds.

## INTRODUCTION

Elucidating routes and levels of migration between populations of a species is essential to understand the forces that shape its evolutionary trajectory (Barrowclough, 1980; Slatkin, 1985). Landscape features—such as rivers, mountain ranges, crop fields, and urban areas—can impact levels of gene flow between populations by determining dispersal rates and routes (McRae, 2006; Cushman *et al.*, 2006) as well as influence the likelihood of successful establishment of immigrants (Wang and Bradburd, 2014; Sexton *et al.*, 2014). Landscape features can also indirectly condition the effect of gene flow by influencing local effective population sizes (Wright, 1949; Slatkin, 1985). Consequently, the landscape, loosely defined as an area with spatially variable biotic and abiotic factors (Holderegger *et al.*, 2010), influences the levels of effective gene flow among populations (Clobert *et al.*, 2012). In this way, the landscape plays a pivotal role in the evolution of species.

In contrast to species that depend almost exclusively on natural dispersal agents, species in heavily human-dominated ecosystems may exploit human activities to maintain gene flow among populations and expand their ranges (Everman and Klawinski, 2013; Fountain *et al.*, 2014). Such species may be capable of maintaining population connectivity over vast geographic ranges (Trakhtenbrot *et al.*, 2005) by overcoming landscape features that would otherwise represent natural barriers. Such species would thus be able to attain dispersal distances that could be orders of magnitude greater than those dependent primarily on natural dispersal agents (Mack and Lonsdale, 2001; Ricciardi, 2007). By facilitating dispersal, humans have the potential to condition the balance between drift and selection (Slatkin, 1985; Lenormand, 2002), introduce genetic variation to local populations (Kolbe *et al.*, 2004), prevent local extinction or favor recolonization (Fountain *et al.*, 2014), and alter the overall genetic constitution of populations (Bataille *et al.*, 2011). Human-aided migration—intentional or unintentional—is particularly prevalent in plants (Hodkinson *et al.*, 2007; Auffret and Cousins, 2013), where it has had major impacts on the distribution of species and stability of communities (Simberloff, 2013 and references therein). Despite our knowledge of both human and natural factors influencing dispersal, there remains a gap in our understanding of the relative influence of each on the distribution of genetic variation among populations of many (if not most) plant species.

A particularly amenable study system to fill this knowledge gap comes from agricultural weed populations. Agricultural weeds experience a highly dynamic landscape characterized by frequent spatial rearrangements and changes in the physical environment (e.g., expansion of agricultural front, increased fragmentation, crop rotation, agricultural chemical use) (Menchari *et al.*, 2007; Meehan *et al.*, 2011). At the same time, natural features such as climate, soil type, and topography likely also play a significant role in structuring populations (Cimalová and Lososová, 2009; Navas, 2012). Under these conditions, human-aided migration may be critical for weedy plant success (Epperson and Clegg, 1986). However, we have limited knowledge of how or if weedy plant populations are able to maintain connectivity through a complex landscape matrix. Addressing this limitation would improve our understanding of the underlying processes governing connectivity of weed populations and also offer practical tools to deal with the economic problems that weeds impose (on the order of 33B USD per year in US agriculture alone; Pimentel *et al.*, 2005).

As a first step into investigating the interplay between natural factors and human activities on structuring genetic diversity in weed populations, we estimate the intensity and extent of migration and evaluate how multiple landscape features influence genetic connectivity of a noxious agricultural weed, *Ipomoea purpurea*. Specifically, we ask the following questions: 1) what is the overall population structure of *I*. *purpurea*, one of the most troublesome weeds in US agriculture (Webster and Nichols, 2012), and 2) which natural and/or human-influenced landscape features—soils, elevation, climate, landcover, crop types, human population density—may act to promote or constrain genetic connectivity between populations of this weed? Answering these questions offers a deeper understanding of the multiplicity of population structure drivers that influence noxious weeds.

## MATERIALS AND METHODS

### Study system

*Ipomoea purpurea*, the common morning glory, is a noxious agricultural weed (Defelice, 2001; Fang *et al.*, 2013) that has a widespread distribution across highly heterogeneous landscapes in the Eastern, South- and Mid-western regions of the United States (Culpepper, 2006; Webster and Nichols, 2012). It is a self-compatible annual bumblebee-pollinated vine and is found primarily in agricultural fields and disturbed areas (Tiffin and Rausher, 1999; Baucom and Mauricio, 2008), as well as cultivated flower gardens and yards (Defelice, 2001). *I*. *purpurea* is one of the most problematic agricultural weeds of southeastern agriculture (Webster and Nichols, 2012), and exhibits variable levels of resistance to the commonly used herbicide glyphosate (Kuester *et al.*, 2015). This species is also a major concern for conservation given its naturalization in multiple regions throughout the world and its aggressiveness as an invasive (Chaney and Baucom, 2012; Fang *et al.*, 2013).

### Data compilation

To capture the plausible effect of both natural and disturbed landscapes on structuring genetic diversity in *I*. *purpurea*, we compiled a diverse set of GIS data for the continental US from a variety of sources (Table S1). These data include human activities (human population density, landcover, planted crops, and roads) as well as natural factors such as elevation, climate (19 variables summarizing central tendencies and variability patterns in temperature and precipitation, and soil characteristics), and soil characteristics (8 variables summarizing the texture, pH, and organic and inorganic content of the top 20cm of soil). Focusing on both sets of data allowed us to assess the relative influence of natural and human effects on structuring *I*. *purpurea*’s populations. We first processed all these data into landscape layers at a common spatial resolution of 10km^2^ and a common spatial extent around the US states with available samples (Fig. 1). This spatial resolution was chosen to maintain a practical balance between scale and analytical manageability given available computational resources. To reduce dimensionality, we opted to perform two separate Principal Component Analyses (PCAs) on the 19 climatic and 8 soil layers, respectively. For all subsequent analyses we kept the resulting first two principal components of each of these analyses, which accounted for over 78% of the variance in each case, and primarily summarized temperature temporal gradients and precipitation seasonality, and soils’ pH, sandiness, and grain size, respectively (Table S2).

**Figure 1.**
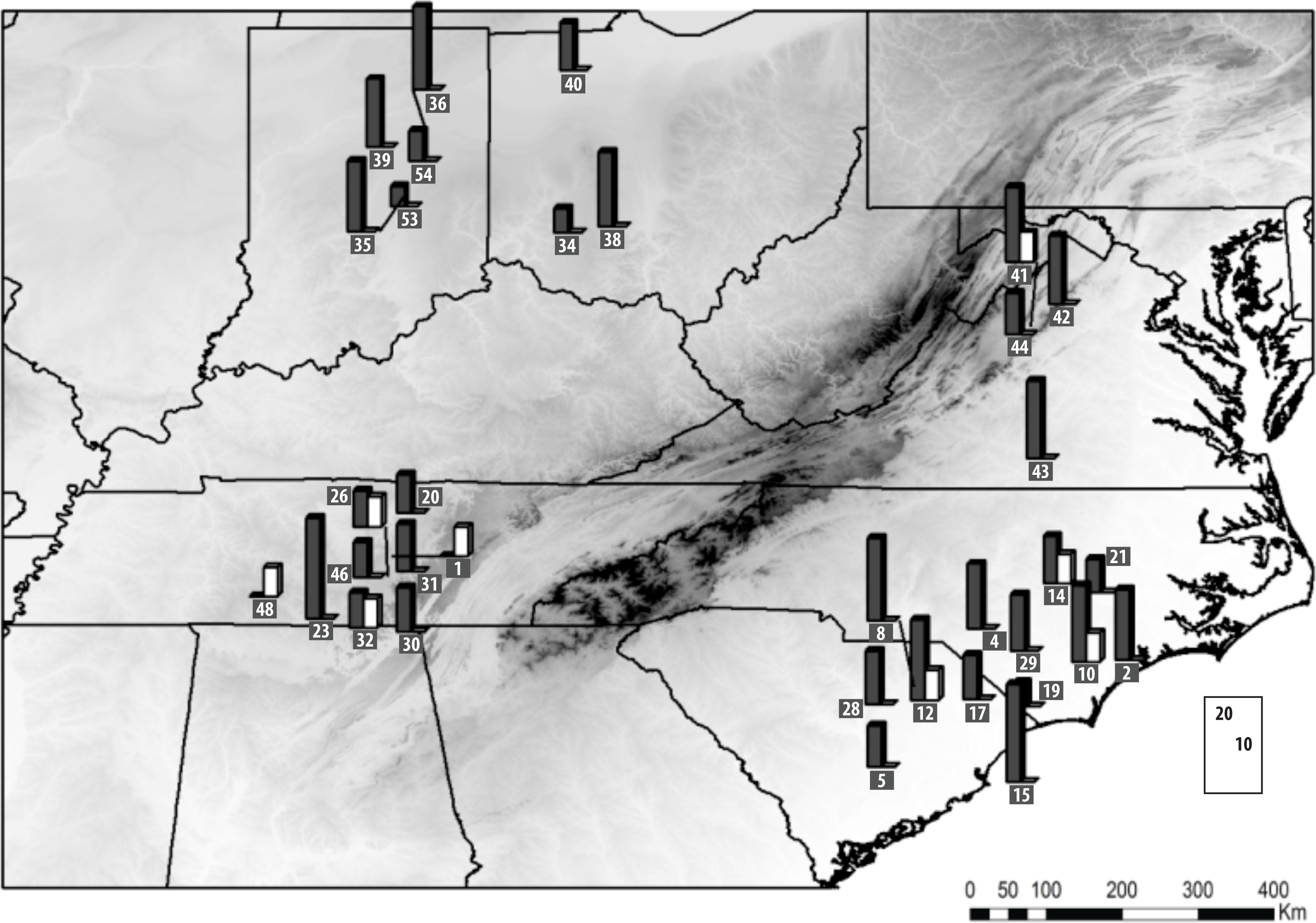
Distribution of *Ipomoea purpurea*’s sampled localities. Sample sizes for both SSR (black bars) and SNP (white bars) datasets are indicated (locality numbers are given in squares). Elevation is provided as background.

We compiled data from a panel of 15 previously optimized microsatellite loci (Molecular Ecology Resources Primer Development Consortium 2013) to examine the genetic connectivity of populations of *I*. *purpurea*. These data (Kuester *et al.*, 2015) encompass a total of 597 individuals from 31 localities (with a minimum of 8 individuals per locality) (Fig. 1; Table S3), collected in 2012 from farms across the range of *I*. *purpurea* in the United Sates (Kuester *et al.*, 2015). In addition, to obtain a more comprehensive representation of the genome of *I*. *purpurea*, we generated a Next Generation Sequencing (NGS) dataset from an additional set of individuals (10 individuals each from 6 localities represented in the SSR dataset, plus 2 additional localities in close geographic proximity to localities in the SSR dataset for a total of 8 populations; Fig. 1).

To generate the NGS dataset, we constructed genome-wide Genotype By Sequencing (GBS) library. The GBS library was developed using 7ng of genomic DNA, extracted from leaf or cotyledon tissue, using SNPsaurus’ (Oregon, USA) nextRAD technology. Samples were first fragmented and then ligated to short adapter and barcode sequences using a partial Nextera reaction (Illumina; California, USA) before being amplified using Phusion^®^ Hot Start Flex DNA Polymerase (New England Biolabs; Massachusetts, USA). The 80 dual-barcoded PCR-amplified samples were pooled and the resulting libraries were purified using AMPure XP beads (Agencourt Bioscience Corporation; Massachusetts, USA) at 0.7x. The purified library was then size selected to 350-800 base pairs and sequenced using two runs of an Illumina NextSeq500 sequencer (Genomics Core Facility, University of Oregon).

The resulting sequences were analytically processed using the SNPsaurus nextRAD pipeline (SNPsaurus, Oregon, USA; Siliceo-Cantero *et al.*, 2016). Specifically, reads of 16 randomly selected individuals (of the 80 sequenced) were combined to create a pseudo-reference genome. This was done after removing loci with read counts above 20,000, which presumably corresponded to repetitive genomic material, and loci with read counts below 100, which presumably corresponded to off-target or read errors. The filtered reads were aligned to each other using BBMap (Bushnell, 2014). All parameters were set to default values with the exception of minimum alignment identity, which was set to 0.93 to identify alleles, as this threshold has been found to work well for non-reference species (SNPsaurus, Oregon, USA). A single read instance was chosen to represent the locus in the pseudo-reference. This resulted in a total of 263,658 loci. All reads from each of the 80 individuals were then aligned to the pseudo-reference using BBMap (Bushnell, 2014) and converted to a vcf genotype table, using Samtools (Li *et al.*, 2009) and bcftools (Li, 2011), after filtering out nucleotides with a quality score of 10 or worse. The resulting vcf table was filtered using vcftools (Danecek *et al.*, 2011) for SNPs with a minimum allele frequency of 0.02, a minimum read depth of 5, and a maximum 15% of missing data. This resulted in 9774 variable regions. Loci with less than 5 high quality base-calls and with more than 20% missing data or an average of less than 20 high quality base calls were also removed using vcftools (Danecek *et al.*, 2011). This resulted in a final panel of quality-vetted 8210 Single Nucleotide Polymorphisms (SNPs) (Fig. S1) that we used in all subsequent analyses.

### Population structure analyses

We first conducted a series of analyses to characterize the overall genetic structure of *I. purpurea* populations. All analyses were run separately for the microsatellite (SSR, hereafter) and SNP datasets given their intrinsic differences and distinct geographic coverage (Fig. 1; Table S3). In addition, we repeated all population structure analyses using just the subset of 6 localities where SSR and SNP datasets are both available. Running separate analyses using these two different markers (referred as SSRc and SNPc, hereafter) allowed us to determine if the differences between marker types was due to differences in sample size (SSR = 24 localities; SNP = 8 localities) or geographic coverage (Fig. 1). Similarly, to determine if the differences uncovered between marker types were due to SNP sequencing or genotyping error, we repeated all population genetic analyses after doing a more stringent SNP quality filtering by removing loci with a genotype quality score below 20, a minimum read depth of 10, or with more than 15% missing data.

First, to characterize population differentiation we estimated F_ST_ using GenAlEx v6.5 (Peakall and Smouse, 2012) (because similar global F_ST_ and R_ST_ estimates were obtained for the SSR dataset, we opted to report F_ST_ values only to allow direct comparisons with the SNP dataset). We then estimated contemporary effective population size for each sampled locality in NeEstimator v2 using the excess heterozygous method (Do *et al.*, 2014). We performed this latter analysis to assess the possibility that differences in local population size underlie differences in genetic variability (Weckworth *et al.*, 2013) and/or promote asymmetric effective migration rate (*Nm*).

In addition, to further examine genetic structure we assessed population admixture and spatial genetic clustering using TESS (Chen *et al.*, 2007). TESS was run using the admixture algorithm and a BYM model (Durand, Jay, *et al.*, 2009) with 10 runs per K value, and without using geographic weights. The TESS model, with the lowest DIC was chosen as the optimal model (Durand, Chen, *et al.*, 2009). K values tested ranged from two to the maximum number of sampled localities. Additionally, following Wang et al. (2009), we complemented these analyses with Analyses of Molecular Variance (AMOVA; Excoffier *et al.*, 1992) run in GenAlEx (Peakall and Smouse, 2012) using 9999 permutation replicates. We ran these AMOVAs either partitioning the variance into regions based on the spatial genetic clusters previously identified by TESS—to quantify the fraction of the genetic variance explained by these clusters, or leaving it ungrouped (i.e., no regions), for comparison.

Additionally, we investigated population connectivity by estimating levels of recent migration between sampled localities through the identification of individuals of mixed ancestry using BayesAss (Wilson and Rannala, 2003). BayesAss is a program that uses individual multilocus genotypes and a Markov Chain Monte Carlo (MCMC) algorithm to probabilistically distinguish between immigrants and long-term native individuals (Wilson and Rannala, 2003). We ran BayesAss for 6 million generations using default parameter settings, and discarded the first two million generations as burn-in (Dyer, 2009). For each marker dataset, we repeated this analysis three times (for a total of 18 million generations) and combined the results from the three replicates for our final inference. Then, using a posterior probability cut-off of 0.75 we assign individuals’ ancestry. We chose this cut-off value as a minimum credibility score to simultaneously maximize sample size and reliability (stringer thresholds show similar differences between marker sets; results not shown). It is important to note that because of computational limits we had to randomly subsample our set of SNPs to 400 SNPs for this analysis. The same subsampled set was used for the full and reduced (SNPc) analyses.

### Landscape genetics analyses

To identify the likely landscape features underlying overall population structure of *I*. *purpurea*, we evaluated the association between landscape features and genetic differentiation based on the full datasets. First, we estimated conditional genetic distances (Dyer *et al.*, 2010) using GeneticStudio (Dyer, 2009). Briefly, conditional genetic distances are measures of pairwise genetic distance derived from population networks, constructed based on the degree of genetic similarity between sampled localities (Dyer and Nason, 2004). They reflect genetic similarity between localities that better capture direct gene flow (i.e., direct migration) as opposed to connectivity driven by step-wise migration through intervening localities (Dyer, 2015). The complexity of the associated conditional genetic network was summarized by their vertex connectivity (White and Harary, 2001), whereas the congruence between networks derived from different marker sets was measured by their structural congruence (a measure of wether the number of congruent edges between networks is greater than expected by chance) (Dyer, 2009).

To assess the association between landscape features and population differentiation, we first converted each landscape layer (climate, crops, elevation, landcover, population density, roads, and soils layers; Table S1) into landscape resistance layers. To do this, each landscape feature in these layers was assigned a resistance value that reflects the difficulty that each feature offers to the movement of gametes or individuals. In contrast to previous studies that typically rely on expert opinion for resistance assignment, we utilized an unbiased statistical optimization to avoid the sensitivity of results to subjective resistance assignment (Spear *et al.*, 2010). Specifically, resistance values were optimized through a genetic algorithm approach (Mitchell, 1996). Briefly, in this search algorithm a population of individuals with traits encoded by unique combinations of model parameters (resistance assignment proposals in our case) is allowed to compete with each other based on the fitness associated with the traits it carries (Peterman *et al.*, 2014). Specifically, in Peterman’s (2014) implementation of this algorithm, which we followed here, individuals’ fitness is estimated by the relative quality of a MLPE.lmm model (Maximum Likelihood Population Effects – Linear Mixture Model). This model evaluates the association between pairwise genetic distance and landscape cumulative resistance between localities, estimated in Circuitscape (Shah and McRae, 2008). Individuals with parameter settings (i.e., resistance assignments) that result in better models, as measured by a Deviance Information Criterion (DIC) score, are preferentially represented in the following generation. Offspring modifications introduced by mutations (i.e., small resistance assignment perturbations) allow for exploration of the parameter space. The algorithm was stopped once 25 generations have passed without significant improvement in fitness.

We implemented Peterman’s (2014) algorithm in R (package ResistanceGA; Peterman, 2014) allowing for the independent optimization of each of our landscape layers. The optimal resistance landscapes identified in this way were then used to run a final univariate MLPE.lmm model to characterize the association between landscape features and conditional genetic distances between localities. Because the roads-associated resistance was not recovered as significant for either marker dataset, we dropped this layer for all subsequent analyses. Finally, to identify the simultaneous contribution of natural and human-driven landscape features to population differentiation in *I*. *purpurea* we ran Multiple Regression on Distance Matrices (MRDM; Legendre *et al.*, 1994). Before running these MRDM models, we standardized all optimized resistance layers to mean of zero and variance of one (Dyer *et al.*, 2010). These final regressions included geographic distance as a null model predictor as well as effective population size and were run in R (package ecodist; Goslee and Urban, 2007) using 10,000 permutations to assess significance. We accounted for multiple testing by applying a false recovery rate correction (Benjamini and Hochberg, 1995) using the function *p.adjust* in R (R Core Development Team, 2016).

These landscape genetic analyses, aimed at identifying the relative influence of natural and human-related landscape features on *I*. *purpurea*’s connectivity, show several differences between SSR and SNP datasets (see below). Nonetheless, we expect that association patterns that are robust between datasets should accurately reflect the impact of landscape features on gene flow, independent of possible biases introduced by marker idiosyncrasies. Therefore, we focus below on the common biological findings between marker types, while also denoting the most relevant differences.

## RESULTS

### Population structure

The initial genetic analyses indicated that *I*. *purpurea* sampled localities were in no violation of Hardy-Weinberg equilibrium (Fig. S1c, d), as evidenced by the small difference between expected and observed heterozygosity (mean He = 0.294±0.014 and 0.250±0.001; mean Ho = 0.291±0.009 and 0.260±0.001, respectively for SSR and SNP datasets). Levels of expected and observed heterozygosity for the SSR dataset were only slightly greater than those estimated for the SNP dataset. Likewise, the estimated mean effective population size per sampled locality was only slightly greater and more variable for the SSR dataset than for the SNP dataset (13.71±5.59, 9.49±0.13, respectively), but in neither case was there salient evidence of a plausible source-sink dynamic, as judged by the similar effective sizes among ^populations. Average FST estimates between datasets were also similar (0.151^ and 0.140, respectively for SSR and SNP datasets; Fig. S2).

Interestingly, we found that estimates of recent ancestry differed between SSR and SNP datasets. The analysis of the SSR dataset indicated that recent migration among localities seems to be more widespread, with only four localities being primarily constituted of native individuals (Fig. 2a). Across localities, on average 73.65% of individuals were inferred to be 1st or 2nd generation immigrants. In comparison, analysis of the SNP dataset showed that most populations seem to have a more limited number of recent immigrants, and that the relatively few inferred immigrants (on average 27.42% of individuals) did not come exclusively from geographically proximate localities (Fig. 2d). Accordingly, SSR and SNP pruned conditional genetic networks (Dyer and Nason, 2004) indicated different underlying patterns of genetic connectivity (structural congruence = 0.108; Fig. 2b,e). While both were fully closed, the SSR-based network was more interconnected (vertex connectivity: 5) than the SNP-based network (vertex connectivity: 0). Further, based on the best TESS models (Fig. S3), widespread admixture was recovered in the SSR dataset (median individual maximum Q-score = 0.51), whereas minimal admixture was identified in the SNP dataset (median individual maximum Q-score = 0.73) (Fig. 2c,f). Finally, grouping individuals according to the corresponding TESS-identified spatial genetic clusters in AMOVA analyses only slightly reduced the variance explained solely by geographic location in both datasets (Table 1).

**Figure 2.**
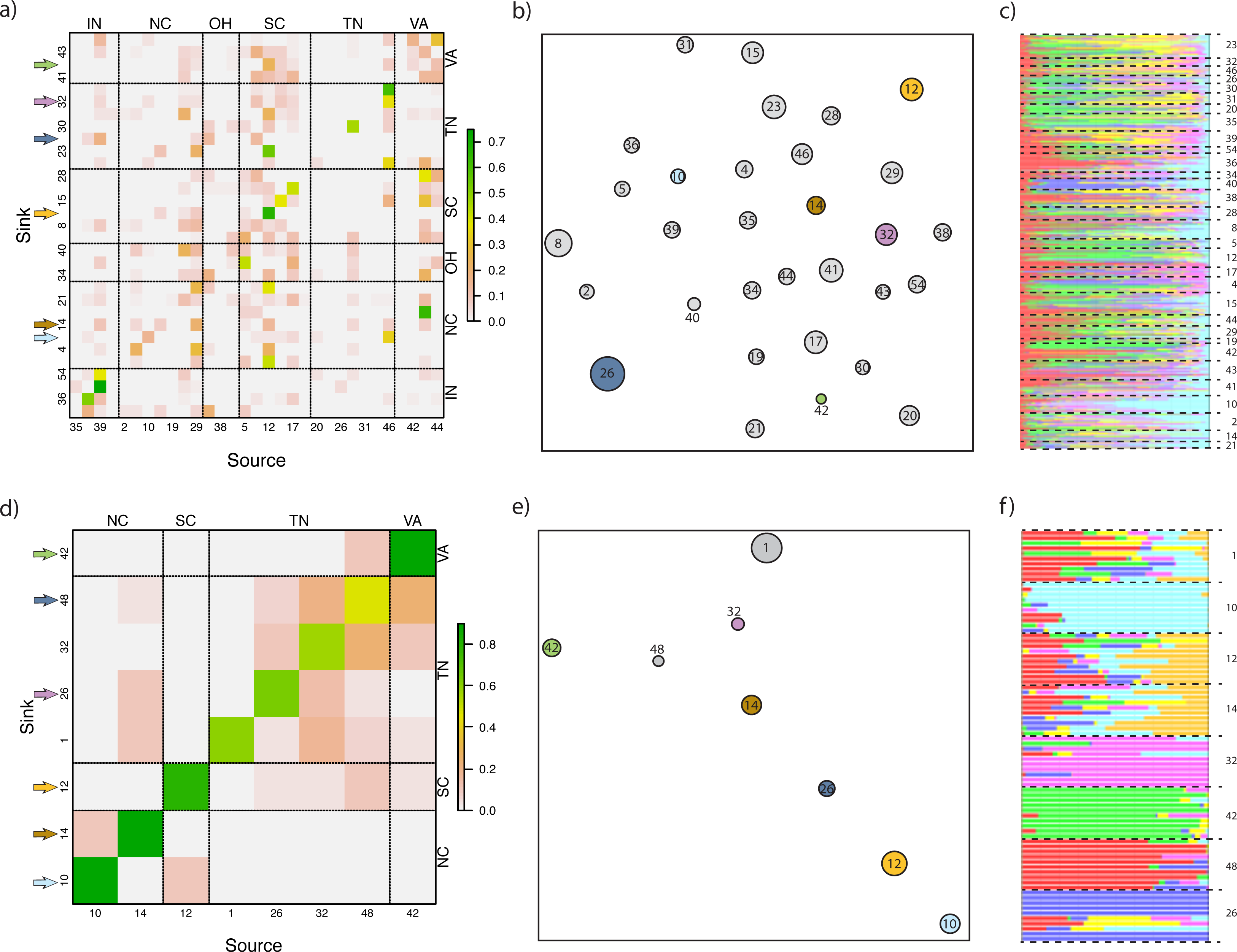
Inferred population connectivity. The estimated origin of individuals for each sampled locality (sink) is depicted according to the locality they were inferred to have originated from (source) (a, d). The color of each cell in these plots depicts the relative proportion of individuals in the sink population that were estimated to be recent immigrants from each locality along the x-axis. Cells on the minor diagonal correspond to the proportion of native individuals. Pruned conditional genetic networks (b, e) and posterior estimates of admixture proportion identified by TESS analysis (c, f) are also displayed. The top row shows SSR-based results, the bottom shows the SNP-based results. Locality numbers follow Fig. 1. Localities shared between SSR and SNP datasets are denoted by colored arrows (for a similar figure based exclusively on these shared localities, see Fig. S4).

**Table 1.**
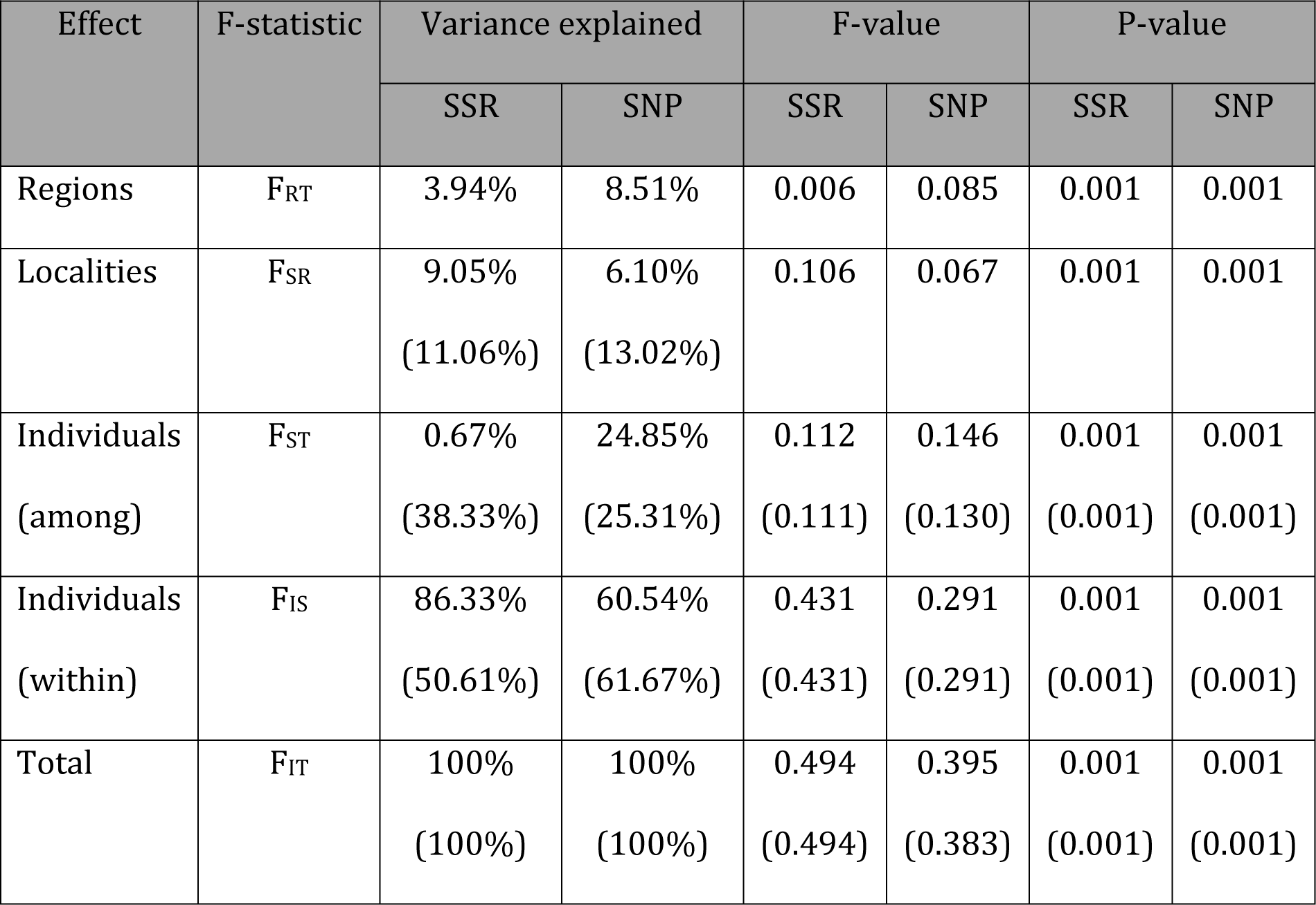
Analysis of Molecular Variance (AMOVA) of SSR and SNP data. The contribution of spatial clusters (regions), localities, and individuals is shown. For comparison, results from an AMOVA analysis with no region category defined are presented in parentheses underneath.

| Effect | F-statistic | Variance explained |  | F-value |  | P-value |  |
| --- | --- | --- | --- | --- | --- | --- | --- |
|  |  | SSR | SNP | SSR | SNP | SSR | SNP |
| Regions | $F_{RT}$ | 3.94% | 8.51% | 0.006 | 0.085 | 0.001 | 0.001 |
| Localities | $F_{SR}$ | 9.05%<br>(11.06%) | 6.10%<br>(13.02%) | 0.106 | 0.067 | 0.001 | 0.001 |
| Individuals<br>(among) | $F_{ST}$ | 0.67%<br>(38.33%) | 24.85%<br>(25.31%) | 0.112<br>(0.111) | 0.146<br>(0.130) | 0.001<br>(0.001) | 0.001<br>(0.001) |
| Individuals<br>(within) | $F_{IS}$ | 86.33%<br>(50.61%) | 60.54%<br>(61.67%) | 0.431<br>(0.431) | 0.291<br>(0.291) | 0.001<br>(0.001) | 0.001<br>(0.001) |
| Total | $F_{IT}$ | 100%<br>(100%) | 100%<br>(100%) | 0.494<br>(0.494) | 0.395<br>(0.383) | 0.001<br>(0.001) | 0.001<br>(0.001) |

Similar to our results from the entire datasets, when we subset the SSR and SNP dataset to the 6 localities in common (SSRc and SNPc datasets), we found no major differences in genetic estimates between the SSRc and SNPc datasets (Table S4), and we again identified differences in the underlying population structure (Fig. S4). Specifically, the SNPc dataset was characterized by a smaller percentage of recent immigrants (28.25%) than the SSRc dataset (44.93%) (Fig. S4a,d), and the corresponding genetic networks were also different from each other (structural congruence = 0.002)—with the SSR-based network being more connected (vertex connectivity = 2) than the SNPc-based network (vertex connectivity = 0) (Fig. S4b,e). Finally, as for the full data, a more admixed genetic composition of individuals was recovered in the SSRc dataset (median individual maximum Q-score = 0.71) than in the SNPc dataset (median individual maximum Q-score = 0.85) (Fig. S4c,f). Also, further confirming the limited spatial structure in this species, using TESS-identified spatial genetic clusters as regions in AMOVA analyses barely reduced the variance explained solely by geographic location when compared to a null model with no regions assigned (Tables S5).

Similarly, when using a more stringently filtered SNP dataset, which comprised 5811 SNPs, we found that this reduced dataset produced highly similar results to the original SNP dataset (percentage of recent immigrants = 25.94%, genetic network vertex connectivity = 0, average individual maximum Q-score = 0.71, and percentage explained by TESS-groupings = 7.31%). Hence, these results using a more rigorous SNP dataset further support the differences in population structure inferences between SSR and SNP data.

### Landscape genetics

Both the SNP and SSR datasets provide evidence that human-impacted landscapes play an important role in shaping genetic connectivity in *I. purpurea*. In both sets of MLPE.lmm models, null (geographic distance), natural (climate, elevation, and soils), and human-related landscapes (landcover and human population density) were identified as significant (p<=0.05) or marginally significant (0.05<p<=0.1) predictors of geonetic differentiation between localities. Interestingly, the variables with the greatest association coefficient and lowest AICc value in both the SSR and SNP models were human-related variables (landcover and human population density, respectively; Table 2). However, when considering all variables together in a multivariate manner—while accounting for geographic distance—human population density, local effective population size, and different aspects of climate were the only variables that remained as significant or marginally significant predictors of genetic differentiation across both SSR and SNP datasets (Table 2). In contrast, elevation and soil were identified as significant or marginally significant predictors only in the SNP dataset. Both multivariate regressions also differed in the proportion of the variance explained (MRDM R^2^ for SSR and SNP dataset were 0.109 (F_1,29_ = 3.654, p-val. = 0.063) and 0.532 (F_1,6_ = 1.932, p-val. = 0.113), respectively).

**Table 2.**
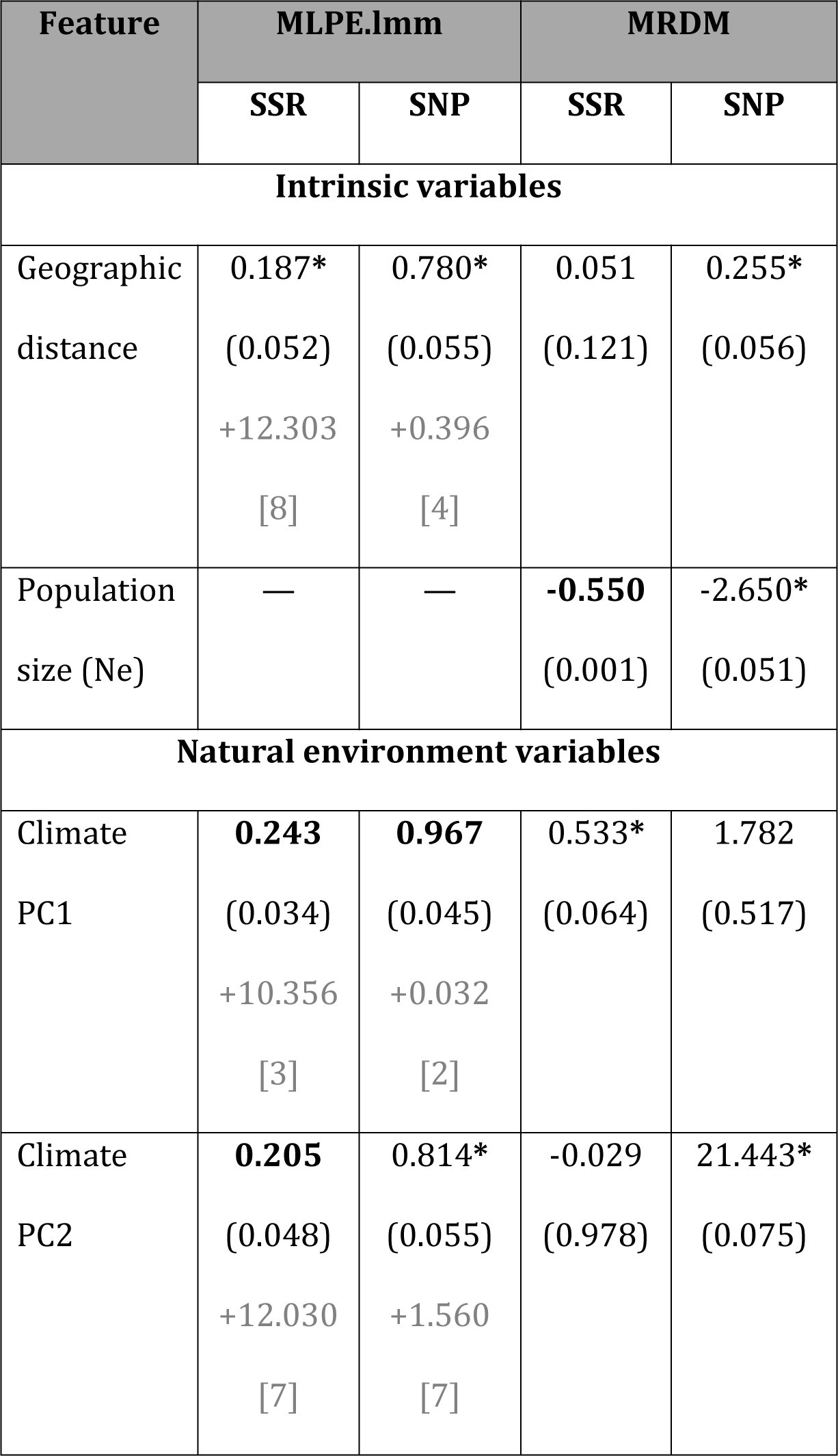
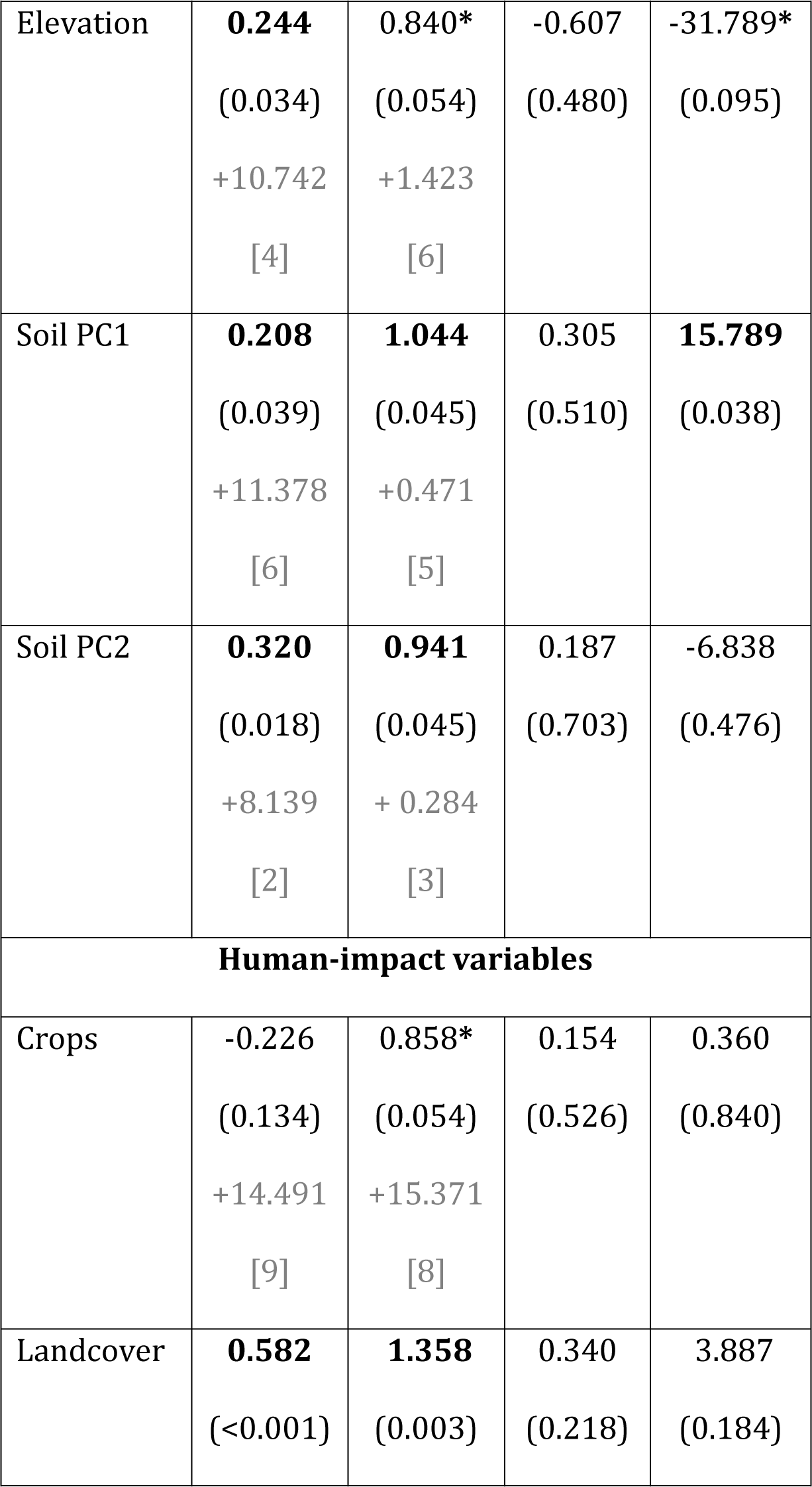
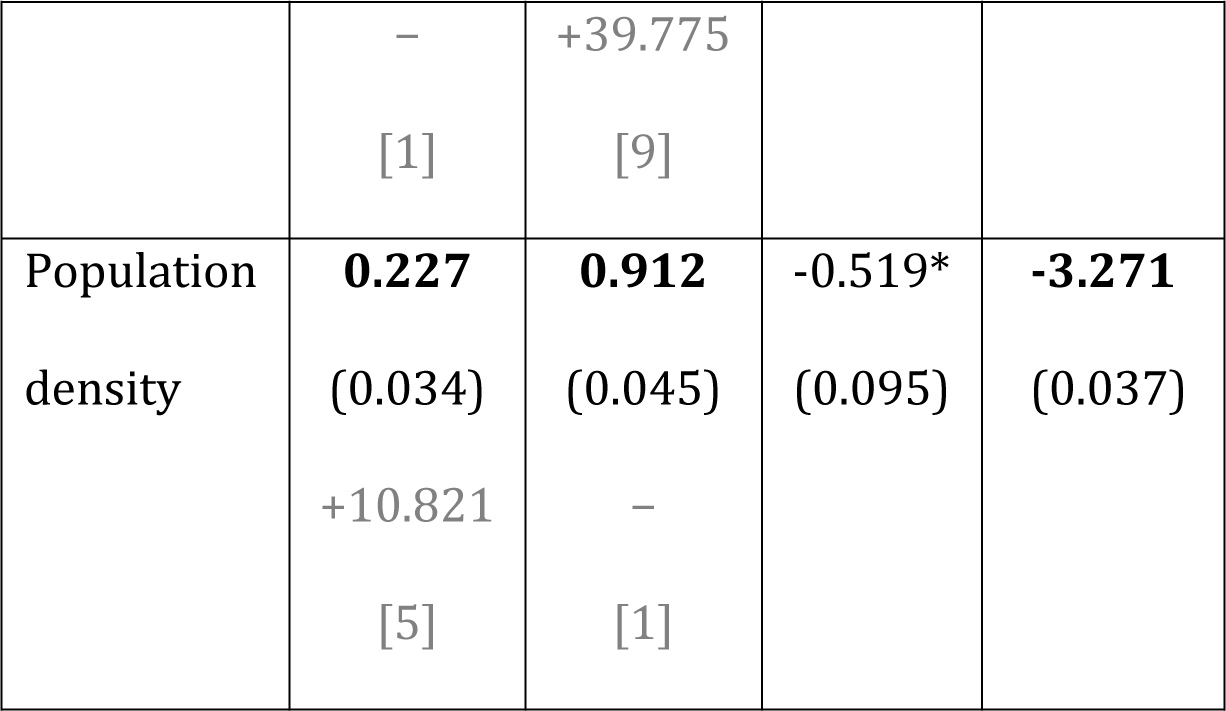
Summary of landscape genetics models. Model coefficients are reported followed by associated p-value (in parenthesis) and, for MLPE.lmm models, followed by AICc difference and ranking (in square brackets). Significant coefficients are in bold, marginally significant coefficients are marked with an asterisk.

In summary, across datasets, results indicated that human-population-density resistance was robustly associated with differentiation among *I*. *purpurea*’s populations, with sparsely to moderately populated areas identified as more conducive areas for migration and potential corridors available between all regions (Fig. S5b). In contrast, climatic variables produced potential barriers to gene flow, with temperature temporal gradients isolating the northernmost localities from the rest in the SSR dataset, and precipitation seasonality isolating the eastern and western localities in the SNP dataset (Fig. S5a). Finally, local effective population size was also a significant predictor in both datasets, with population size inversely associated with genetic differentiation (Table 2).

## DISCUSSION

Our results reveal that broadly distributed populations of *I*. *purpurea* are not genetically isolated from each other. They also suggest the existence of long-distance and putatively human-mediated migration between localities. At the scale of our analyses, the regional agricultural matrix does not seem to have an overarching impact on population connectivity in this species, despite *I*. *purpurea*’s tight link to agricultural fields. Instead, genetic connectivity in this species seems to be primarily influenced by climate and human population density. The effective population size (Ne) also appears to influence population connectivity in this species, which suggests a plausible additional effect of genetic drift on the effectiveness of gene flow (Weckworth *et al.*, 2013). Taken together, these results highlight the significant interplay between human-driven and natural landscapes in structuring *I. purpurea* populations.

### Population connectivity patterns

Despite *I*. *purpurea*’s expected low natural seed dispersal, due to heavy, gravity dispersed seeds, and large and patchy distribution, we found evidence of limited genetic differentiation through the species’ range and overall weak geographic structure. In fact, population differentiation was uneven (pairwise ^FST values ranged from 0.02 to 0.24), and only partially dependent on the^ geographic distance between populations. Further, genetic networks for both datasets were fully closed, suggesting the existence of direct or indirect gene flow among all sampled populations. In line with this finding, admixed individuals were present in all sampled localities (although levels of admixture vary for SSR and SNP datasets), and several instances of recent short- and long-distance migration were recovered. Nevertheless, the evidence of interconnectedness we uncovered is likely also influenced by the shared evolutionary history of populations and, thus, shared ancestral genetic variation likely confounds our estimates of genetic differentiation (Marko and Hart, 2011). Given the relatively recent invasion of the US by *I*. *purpurea* and its inclusion in horticultural trade (Fang *et al.*, 2013), it is possible that historical connectivity between populations due to human transport maintained gene flow between populations after its introduction into the US (Mack, 1991). Alternatively, recurrent colonization from few genetically similar sources might instead have resulted in partial homogenization of otherwise isolated populations (Dlugosch and Parker, 2008). To disentangle the relative contribution of recent gene flow from that of historical patterns of population connectivity, empirical estimates of inter-population migration (e.g., through the use of pollen traps) or direct measurements of gene flow (e.g., paternity analysis) would be needed.

Our estimates of differentiation as well as heterozygosity and allelic richness differ from other agricultural weedy species. Specifically, our estimates suggest that *I*. *purpurea* has lower genetic diversity (as measured by allelic richness and/or heterozygosity) than primarily outcrossing annual weeds (e.g., ragweed (Martin *et al.*, 2016) and black-grass (Menchari *et al.*, 2007)), including many broadly distributed invasives (Dlugosch and Parker, 2008). This finding is potentially explained by both differences in reproduction system, as outcrossing plants often show greater genetic diversity than plants with mixed or primarily selfing reproduction (Hamrick and Godt, 1996), and the recent population bottlenecks likely experienced by *I*. *purpurea* (Kuester *et al.*, 2016). Genetic differentiation found in this study (as measured by F_ST_), on the other hand, was relatively low for a broadly distributed gravity-dispersed plant (Hamrick and Godt, 1996). This contrasts with many broadly distributed weeds, which show moderate to high F_ST_ values (0.15 - 0.48) likely due to dispersal limitations over distances above those commonly allowed by natural dispersal agents (Schmidt *et al.*, 2009; Treier and Müller-Schärer, 2011). This is the case, for example, of invasive weeds such as weedy *Silene* (Barluenga *et al.*, 2011) that rely on specialist pollinators or seed dispersers that might not be present in the invaded range. Considering the natural history of *I*. *purpurea*—heavy seeds, bumblebee pollination (Osborne *et al.*, 1999; Schulke and Waser, 2001), and strong agricultural and horticultural ties (Defelice, 2001), this finding suggests that human-aided dispersal presumably contributes to maintain connectivity in this species.

### Landscape features influencing population connectivity

Our assessment of the association between population differentiation and landscape resistance identified several predictors of population connectivity. Of the landscape features examined, climate and human population density are the only robust predictors, suggesting a role for both human activities and climatic barriers in shaping population connectivity in this species. Specifically, our results suggest that while climatic variables provide some resistance to migration, scarcely to moderately populated areas (i.e., those corresponding with rural areas) offer potential corridors between all sampled regions. The existence of climatic dispersal barriers presumably arises in response to physiological preferences or local adaptations of the species (Cimalová and Lososová, 2009); humans seem to oppose these natural limitations and help *I*. *purpurea* overcome climatic barriers (e.g., by allowing dispersal between northern and southern populations that are separated by areas of high temperature-related resistance to gene flow). Indeed, human-mediated migration is expected to be particularly prevalent among wild populations of species with commercial value such as *I*. *purpurea* (Mack, 1991; Defelice, 2001).

Population connectivity of species in human-dominated environments is potentially also affected by human-induced changes in population size (Méndez *et al.*, 2014). Specifically, by influencing population size, anthropogenic activities condition the effectiveness of migration as it depends on migration relative rate (*Nm*) (Wright, 1931). Furthermore, population size differences influence overall dispersal spatial dynamics because the number and direction of migrants depend on local population sizes (e.g., proportionately greater number of migrants move from densely to sparsely populated areas than vice versa; Lenormand, 2002). In line with these expectations, we found that effective population size is also a significant predictor of population differentiation in this species. Considering the prevalence of weed management practices (e.g., tillage, herbicide application) and their effect on weed population sizes (Kuester *et al.*, 2016), it is likely that these anthropogenic activities play a significant role in controlling the rate of population differentiation in weeds. Thus, all evidence suggests a predominant role of human activities in shaping *I*. *purpurea*’s current genetic structure.

Relatively few studies have explored how landscape features impact population connectivity in weeds at large spatial scales, making it hard to evaluate how distinctive *I*. *purpurea*’s response to climatic and anthropogenic factors may be from that of other weeds. Yet, simulations modeling short-distance dispersal based on spatial distances and landscape configuration have identified dispersal capabilities and landscape use (including availability of disturbed habitats and distribution of crop types) to be the most prevalent determinants of local level connectivity in several weed systems (Woolcock and Cousens, 2000; Fénart *et al.*, 2007; Will and Tackenberg, 2008). For instance, using a ~10km^2^ aerial photograph to inform a spatial mechanistic model of Canadian horseweed’s interfield dispersal, Dauer et al. (Dauer *et al.*, 2009) showed that distribution of suitable habitat primarily determined the rate and extent of this weed’s dispersal at this spatial scale. In agreement with these predictions, empirical data show that local dispersal in mountain pasture weed is heavily influenced by the spatial distribution of human-dominated landscapes and the opportunities for interfield contamination (Treier and Müller-Schärer, 2011). Yet, while neither landcover nor crop types distribution were identified as significant predictors in our multivariate analyses, we cannot rule out the possibility of high local gene flow at the scale of contiguous agricultural fields mediated by these landscape features given the scale of our analyses.

### Marker-specific inferences

Our results identified interesting differences between marker types in terms of the inferred population structure of *I*. *purpurea*. The differences uncovered between datasets are at first glance unexpected as all loci in a species’ genome evolve under a common evolutionary history (Payseur and Cutter, 2006). Nevertheless, differences between marker types have been similarly observed in other studies (Dixon *et al.*, 2011; Martin *et al.*, 2016). For ^example, several studies have recovered FST estimate differences when using^ SNP or SSR loci on the same set of samples (Coates *et al.*, 2009; Gärke *et al.*, 2012), presumably due to mutation rates and genomic representation differences of SSR and SNP loci (Payseur and Cutter, 2006; Coates *et al.*, 2009). Here, the population structure differences we uncovered may likewise be related to marker-specific rates of mutation, drift and/or marker-specific biases—such as greater ascertainment bias on SSR data (Väli *et al.*, 2008; Defaveri *et al.*, 2013)—and not likely caused by the different SSR and SNP dataset sampling. Because of these intrinsic marker differences, both markers could provide complementary information (Payseur and Cutter, 2006). While our identification of robust environmental predictors of genetic differentiation for SSR and SNP datasets—despite differences in underlying population genetic patterns—is encouraging, further work is needed to reconcile traditional landscape genetics studies based on a few highly variable markers with increasing landscape genomics studies based on thousands of SNPs.

### Conclusions

While our results suggest that *I*. *purpurea* experiences moderate inter-population connectedness and potential long-distance dispersal, we cannot rule out the influence of historical relatedness on the observed genetic patterns. To determine the amount of current-day gene flow that occurs between populations, direct estimates of gene flow need to be examined. Regardless, the limited population structure recovered as well as the identification of human population density as a significant predictor of population differentiation calls attention to the need for investigating the possible impact of human-mediated gene flow on the evolutionary path of this species—including its response to selection and the likelihood of further naturalization. In particular, it remains to be investigated whether the pattern of maintained connectivity we identify here could facilitate the success of this weed (e.g., by introducing relevant genetic variants; Kolbe *et al.*, 2004) or reduce the fitness of local populations (Keller *et al.*, 2000). What seems clear from this study is that human-aided migration presumably is an important component of gene flow between populations, which may counter the isolating effects of natural environmental barriers and genetic drift.

## ACKNOWLEDGEMENTS

We thank Adam Kuester for seed collecting and for contributing valuable data for this study. We also thank Ariana Wilson, Eva Fall, and Dan York for tissue collection. This research was funded by USDA NIFA grants 04180 and 07191to R.S.B.

## DATA ARCHIVING

All data generated is in the process of being archived in Dryad. The corresponding doi would be made available upon acceptance.

